# A Genetically Based Latitudinal Gradient in Aggressiveness of Root-Knot Nematode Populations on Tomato with the *Mi-1* Resistance Gene

**DOI:** 10.1101/2025.06.18.660456

**Authors:** Damaris Godinez-Vidal, Francisco Franco-Navarro, Scott Edwards, Antoon T. Ploeg, Simon C. Groen

## Abstract

Tomato is a food crop of global importance, with a large proportion of processing tomatoes produced in California. One of the main problems plaguing tomato production is infection by populations of root-knot nematodes (RKNs; *Meloidogyne* spp.) that can evade or suppress (‘break’) resistance mediated by the *Mi-1* gene, which has been introgressed into most processing tomato cultivars. In this study, we evaluate fourteen *Meloidogyne* spp. populations collected from fields across the state of California that can complete their life cycle on *Mi-1* tomato cultivars. One of these populations was identified as *M. javanica* and the others as *M. incognita*. All RKN populations developed and reproduced on *Mi-1* tomato cultivar ‘Celebrity’. Although we did not observe differences in gall index among populations when studied together in greenhouse conditions, significant quantitative variation in reproduction factor values was apparent among them. Several pathogens and parasites display geographical gradients in aggressiveness and we identified a negative correlation between populations’ latitudes-of-origin and reproduction factors. This suggests that populations of these thermophilic RKN species from lower latitudes tended to have evolved higher levels of aggressiveness on *Mi-1* tomato, which may be linked to warmer temperatures throughout the year. Finally, populations with relatively low reproduction factor values still showed significant differences in the phenotypes of the galls they induced. Our results showed that *Meloidogyne* populations evolved genetic variation in aggressiveness along a latitudinal gradient in Californian processing tomato agroecosystems, which may have implications for managing these important crop pests.

## Introduction

Tomato production in California plays a vital role in the state’s agricultural economy and the global food industry. More than 90% of processing tomatoes in the US are grown in California, along with a significant portion of tomatoes for fresh market consumption. The state recently produced an average of 12.7 million tons of processing tomatoes annually (USDA NASS, 2023). The Central Valley—mainly Fresno, Yolo, and San Joaquin counties—has favorable growing conditions for the crop due to its Mediterranean climate, fertile soils, and advanced irrigation systems (Hartz et al., 2008). However, one of the main problems hampering tomato production in California is infection by RKNs, particularly the thermophilic species *M. incognita*, *M. javanica*, and *M. arenaria* (Dávila-Negrón and Dickson, 2013; Ploeg et al., 2023). Annual relative yield losses of vegetable crops caused by *Meloidogyne* spp., in both temperate and tropical countries, have been estimated at around 20%, but reach up to 80% in heavily infested soils (Koenning et al., 1999; Sikora and Fernandez, 2005). Over the years, strategies developed for managing RKNs include the use of crop rotation, resistant crop cultivars, organic amendments, biological control agents, and chemical compounds such as soil fumigants or novel non-fumigant nematicides (Lopez-Perez, 2006). The use of resistant cultivars has been the primary strategy to control infections by these three major RKN species in tomato (Roberts, 1992; Williamson, 1998). Genetically mediated resistance of available commercial processing tomato cultivars is mainly based on introgression of a single dominant gene from the wild relative *Solanum peruvianum*, the *Mi-1* gene, which encodes an intracellular nucleotide-binding domain and leucine-rich repeat (NLR) immune receptor (Gilbert and McGuire, 1956; Kaloshian et al., 1996; Williamson, 1998). This gene’s widespread use resulted in high selection pressure on RKNs and pushed multiple *Meloidogyne* populations towards evolution of mechanisms for evading or suppressing (‘breaking’) *Mi-1*-mediated resistance. Indeed, growers are now observing RKN infections of their previously resistant tomato crops in increasing numbers (Ploeg et al., 2023).

*Meloidogyne* populations infecting and reproducing in *Mi-1* tomato cultivars have been reported from tomato crops worldwide (Castagnone-Sereno et al., 1993; Okamoto and Mitsui, 1974; Tzortzakakis et al., 2005). The evolution of resistance-breaking phenotypes in *Meloidogyne* populations has been associated with constant exposure to *Mi-1* tomato cultivars (Meher et al., 2009), but populations with such phenotypes have even been isolated from fields with no history of tomato cultivation (Kaloshian et al., 1996; Tzortzakakis et al., 2005; Hajihassani et al., 2022). Recently, Ploeg et al. (2023) reported genetic variation in aggressiveness among five *Meloidogyne* populations from field-grown tomatoes in different counties of California. Four of these populations were identified as *M. incognita* and one as *M. javanica*. In this study, we compare aggressiveness of these five *Meloidogyne* populations to an additional nine *M. incognita* populations, all of them collected from fields planted with *Mi-1* tomato cultivars across the state. Since geographical variation in aggressiveness has evolved along environmental gradients for several pathogens and parasites (Hargreaves 2024; Laine, 2008; Pariaud et al., 2009; Springer, 2007), we further aimed to study if correlations exist between the aggressiveness of these fourteen *Meloidogyne* populations on *Mi-1* tomato and their geographical origins.

## Materials and Methods

### Origins and propagation of *Meloidogyne* populations

*Meloidogyne* populations were obtained from processing tomato cultivars carrying the *Mi-1* gene across California (Table 1). The collection of thirteen of these populations and an initial characterization of the aggressiveness of five of them has been described previously (Ploeg et al., 2023). *Meloidogyne* populations were maintained for multiplication on plants of *S. lycopersicum* cultivar ‘Celebrity’, a hybrid harboring the *Mi-1* gene (Bayer/Seminis, St. Louis, MO), but on which these populations are virulent. Roots were transferred to new pots to continue population multiplication. As a control, we used a “standard” avirulent population P77R3 (Project 77/Race 3; also known as VW6), to which the Celebrity cultivar shows resistance. P77R3 was maintained on *S. lycopersicum* cv. Cherry (Osborne Seed). Plants were grown under greenhouse conditions at soil temperatures ranging from 21 °C to 26 °C and watered and fertilized daily under an automated irrigation system.

**Table 1.** Source (county and field-grown tomato cultivar) of fourteen *Meloidogyne* populations virulent on *Mi-1* tomato (modified from Ploeg et al., 2023).

| ID | County of origin | Tomato cultivar |
| --- | --- | --- |
| 1 | Yolo | HEINZ 5508 |
| 2 | San Joaquin | SUN6366 |
| 3 | Yolo | HEINZ 5508 |
| 4<sup>1</sup> | Kern | CXD 187 |
| 7 | Kern | HEINZ 5508 |
| 8 | Fresno | UG19406 |
| 11 | Yolo | HEINZ 5508 |
| 12 | Yolo | HEINZ 5508 |
| 15 | Sutter | H1310 |
| 17 | Yolo | HEINZ 5508 |
| 20 | Colusa | HEINZ 5508 |
| 21 | Colusa | N6416 |
| 22 | Sutter | H1310 |
| 26<sup>2</sup> | Merced | HEINZ 5508 |

<sup>1</sup>Identified as *M. javanica*.
<sup>2</sup>Identified as *M. incognita* by observations of perineal patterns and PCR-based identification.

### Identification of *Meloidogyne* populations

Except for one population, the *Meloidogyne* populations have been identified previously: one as *M. javanica* and another twelve as *M. incognita* (Ploeg et al., 2023). The one population of unknown identity was propagated following the approaches described above. To collect females for morphological identification (Eisenback, 1985), infected tomato roots were cleared with 10% bleach for 5 min and stained with 1× acid fuchsin (Sigma-Aldrich) solution (1.75 g acid fuchsin, 125 mL acetic acid, and 375 mL H_2_O). Perineal patterns were cut and placed in 60% glycerol on slides and examined using a Leica optical Microscope, objective 10×/0.5 Plan H, with a Nikon DS Camera Head and Nikon NIS Elements Imaging Software to process images. Morphological observations were confirmed via PCR-based identification. Genomic DNA (gDNA) was extracted from females of population #26 using the CTAB method (Godinez-Vidal and Groen, 2025). One hundred nanograms of gDNA were used to amplify regions of genes encoding NADH dehydrogenase subunit 5 (*NAD5*; Janssen et al., 2016), an esophageal gland protein (*SEC1*; Tesarová et al., 2003), and cytochrome oxidase subunit II (*COII*; Table 2; Powers and Harris, 1993). The PCR reaction contained 5 µL of Apex 2× TaqRed Master Mix, 0.5 µL of each primer (10 pmol), and 3 µL of H_2_O. The PCR program started with 94°C for 5 min, followed by 35 cycles of 94°C for 30 sec, 50°C for 30 sec, and 72°C for 1 min, as well as a final extension at 72°C for 5 min. Ten μL of the amplicon suspensions were run on a 1% agarose gel in TAE buffer at 120 V for 30 min. For the positive control DNA reference, we used DNA from population P77R3 / VW6, previously identified as *M. incognita* based on whole-genome resequencing data (Szitenberg et al., 2017).

**Table 2.**
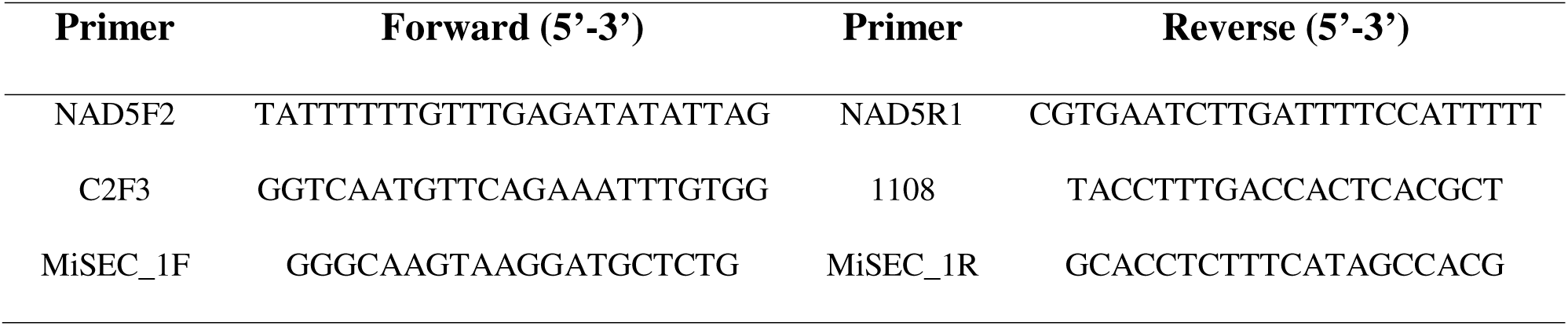
Primers used for molecular identification of *Meloidogyne* population #26.

### Plant preparation and nematode inoculation

Seeds of tomato cv. Celebrity were planted in peat moss (Sunshine Mix 5, Sungro) on nursery seedling trays. After three weeks, the seedlings were transplanted to 1-L pots containing UC mix soil (90% sand, 10% organic matter). One week after transplant, plants were fertilized with Osmocote (Miracle-Gro Co.) and kept under greenhouse conditions. Eggs from each *Meloidogyne* population, including population P77R3 / VW6, were extracted from infected roots (Godinez-Vidal et al., 2024), collected, and quantified. In each 1-L pot, three 5-cm-deep holes were made around the stem of the tomato plant and 80,000 eggs were inoculated per *Meloidogyne* population. After inoculation, pots were lightly watered by hand. Pots inoculated with the same RKN population were then placed together, slightly separated from pots inoculated with other RKN populations, on a bench under the same greenhouse conditions as described above. Each of the 15 populations were tested in 3 replicate pots per trial and the greenhouse trial was replicated twice. One week after inoculation, plants were watered and fertilized (17-6-10 controlled-release fertilizer, Scotts-Sierra Horticultural Products Co) via water drips connected to an automated drip system to avoid contamination from splashing. Eight weeks after inoculation, plants were removed from the pots, and roots rinsed with water to remove excess soil. Roots were indexed for root galling severity using a 0-10 Gall Index (GI) scale, where 0 indicates no observable galling and 10 indicates that 100% of the root system was galled. Once rated, roots underwent egg extraction (Godinez-Vidal et al., 2024). Eggs collected were quantified using a dissecting stereo microscope at 20× magnification. RKN reproduction was expressed as the reproduction factor (RF), where Pf/Pi = number of eggs per root system at harvest/number of eggs inoculated.

### Statistical analysis

Differences among *Meloidogyne* populations on tomato cv. Celebrity were analyzed statistically using GraphPad Prism Software (Version 10.4.2). The data from the two greenhouse trials were analyzed together. Two-way analysis of variance (ANOVA) was used to test the main effects of trial and inoculation with different *Meloidogyne* populations as well as the interaction between these main effects for the RF values. Between-population differences in RF values were determined using Tukey’s HSD post-hoc tests. Differences in GI were analyzed using non-parametric Kruskal-Wallis tests. Pearson’s product-moment correlation coefficients were estimated to determine relationships between the across-trial means of GI and RF values of different *Meloidogyne* populations and the latitudes of their origins. Gall length and area were measured using Fiji-ImageJ2 Software (Version 2.16.0/1.54p). Differences in gall size measurements were analyzed using non-parametric Kruskal-Wallis tests.

## Results

### *Meloidogyne* population #26 belongs to *M. incognita*

Perineal patterns of adult female RKNs from *Meloidogyne* population #26 corresponded to *M. incognita*: the high, squarish dorsal arch with wavy, forked striae and an oval shape was consistent with characteristics reported for this species; we did not observe distinct lateral fields and the striae in the ventral region were relatively straight (Fig. 1A-C). PCR-based identification confirmed the species identified from these morphological observations. Amplifying regions of the *NAD5* and *SEC1* genes from this population, resulted in amplicons of 600 and 500 bp, respectively (Fig. 1D), which would be expected for *M. incognita*. Furthermore, we obtained an expected partial amplicon of 1.2 kb for the *COII* gene. We conclude that this population belongs to *M. incognita*.

**Fig. 1.**
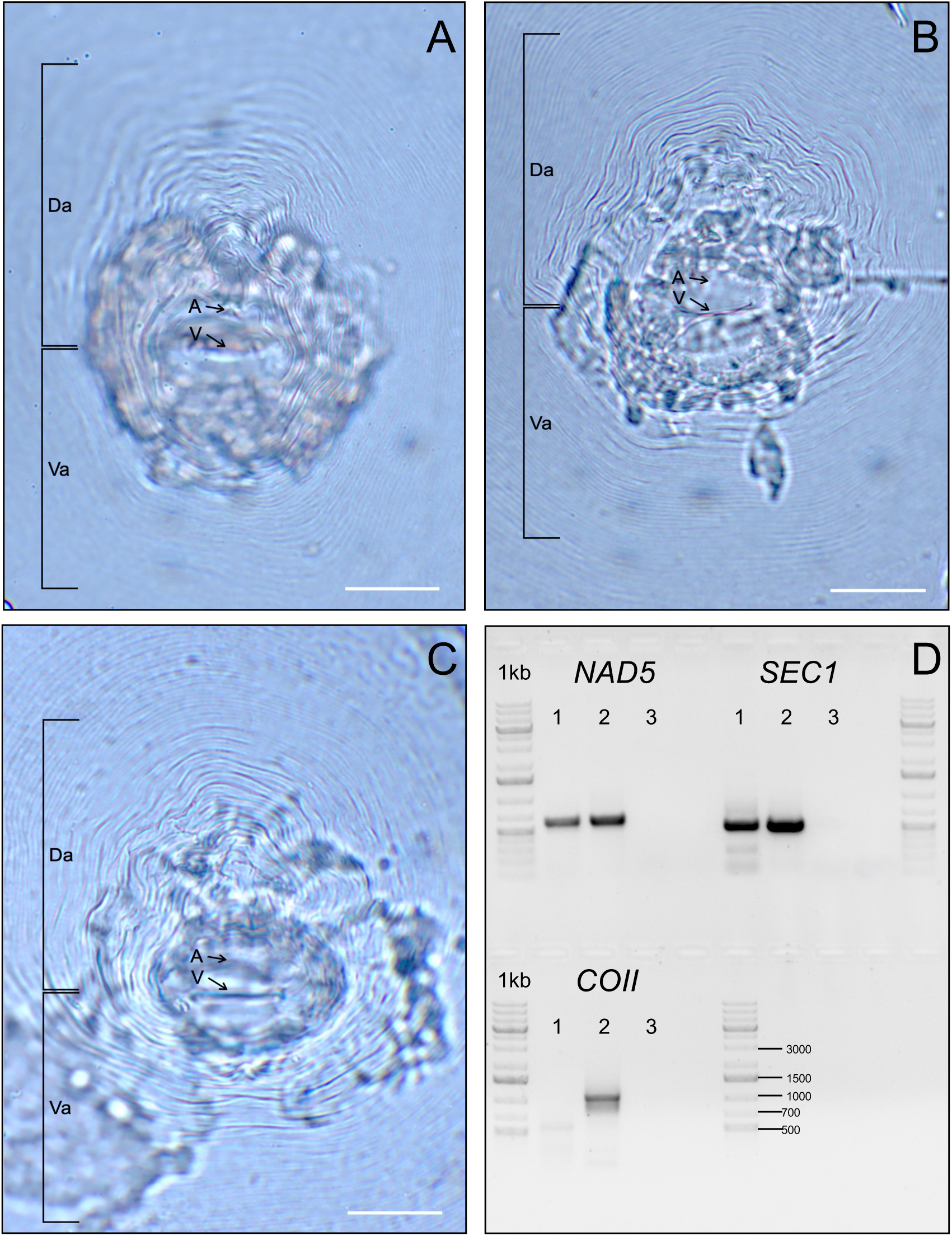
Identification of *Meloidogyne* population #26 as *M. incognita*. A-C. Representative perineal patterns of *Meloidogyne* population #26. Da = dorsal arch, Va = ventral arch, A = anus, V = vulva. D. PCR-based DNA amplification of regions of the *NAD5*, *SEC1*, and *COII* genes from nematodes of population #26, shown alongside 1-kb ladders. Lane 1 contains amplified DNA from *M. incognita* population P77R3 / VW6 (previously confirmed to be *M. incognita*), Lane 2 contains amplified DNA from population #26, and Lane 3 represents a negative control without DNA.

### A genetically based latitudinal gradient in aggressiveness of *Meloidogyne* populations on ***Mi-1* tomato**

All *Meloidogyne* populations were able to reproduce on *Mi-1* tomato cv. Celebrity plants and caused root GIs >7; no significant differences in GI values were observed among the populations (Fig. 2A). In contrast, we observed significant differences in RF values among the populations (P<0.001; Fig. 2B). Statistical analysis divided the populations in three groups: one containing populations with RF values >10 (#4, 17, and 20), a second group with RF values >6 (#1, 7, 8, and 12), and a third group with RF values >2 (#2, 3, 11, 15, 21, 22, and 26). The avirulent P77R3 / VW6 population behaved as expected, with very minor root galling and an average RF value of 0.2.

**Fig. 2.**
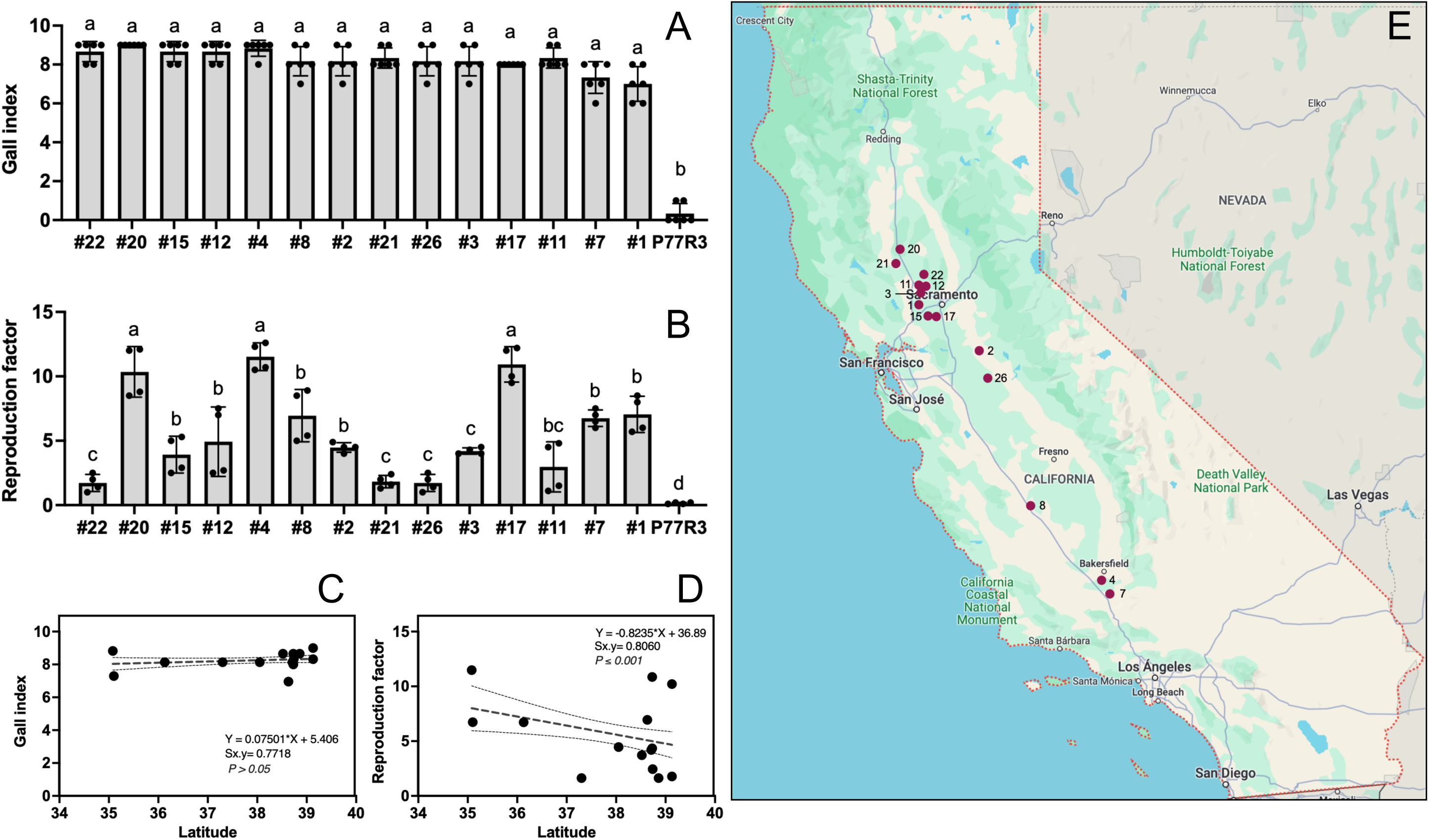
A genetically based latitudinal gradient in aggressiveness of *Meloidogyne* populations on *Mi-1* tomato. A. *Meloidogyne* populations virulent on *Mi-1* tomato did not differ significantly in their gall indices on *Mi-1* tomato cv. Celebrity across two greenhouse trials (Kruskal-Wallis tests). *M. incognita* population P77R3 / VW6 is avirulent on *Mi-1* tomato and was included as a control. B. *Meloidogyne* populations virulent on *Mi-1* tomato differed significantly in their reproduction factor values on *Mi-1* tomato cv. Celebrity across two greenhouse trials (two-way ANOVA with Tukey’s HSD post-hoc tests). Letters above treatment groups in A and B indicate statistically significant differences. C. Latitudes-of-origin and gall indices are not significantly correlated between *Meloidogyne* populations (Pearson’s product-moment correlation, P>0.05). D. Latitudes-of-origin and reproduction factor values are significantly correlated between *Meloidogyne* populations (Pearson’s product-moment correlation, P<0.001). Data shown are means across trials in A-D as well as standard errors of the mean in A and B. E. Source environments of fourteen *Meloidogyne* populations used in this study that had been collected throughout California from *Mi-1* processing tomato cultivars.

Next, we determined whether the geographical source locations of the populations were related to their GI and RF values by estimating Pearson correlation coefficients between these two factors and populations’ source environments. We particularly focused on latitude since the collection locations of the populations were distributed along a latitudinal axis across California. As expected, no correlation was observed between the latitudes-of-origin and GI values among populations (Fig. 2C). However, a negative correlation was identified between latitudes-of-origin and RF values (P<0.001; Fig. 2D), suggesting that populations that occur at lower latitudes tended to have evolved higher rates of reproduction and aggressiveness on *Mi-1* tomato than populations occurring at higher latitudes (Fig. 2E).

### Galling and root damage induced by *Meloidogyne* populations on *Mi-1* tomato

When observing a high GI value, a high RF value is usually expected. However, our results showed high GI values combined with relatively low RF values for some of the *Meloidogyne* populations: #2, 3, 11, 15, 21, 22, and 26. In addition to their lower RF values, these populations showed a galling phenotype distinct from those observed for the other *Meloidogyne* populations. These populations induced large galls rather than inducing a combination of several small galls as observed for the populations with higher RF values (Fig. 3A). To make these observations quantitative, we measured the length and area of the galls induced by populations #2, 3, 11, 15, 21, 22, and 26. We observed significant differences in gall length among these populations (P<0.001). Populations #2, 15, 22, and 26 induced the longest galls, followed by populations #3 and 11, and, finally, population #21 (Fig. 3B). Regardless of the differences in the length of the induced galls, no significant differences among these populations were observed in the area of the galls induced (Fig. 3C).

**Fig. 3.**
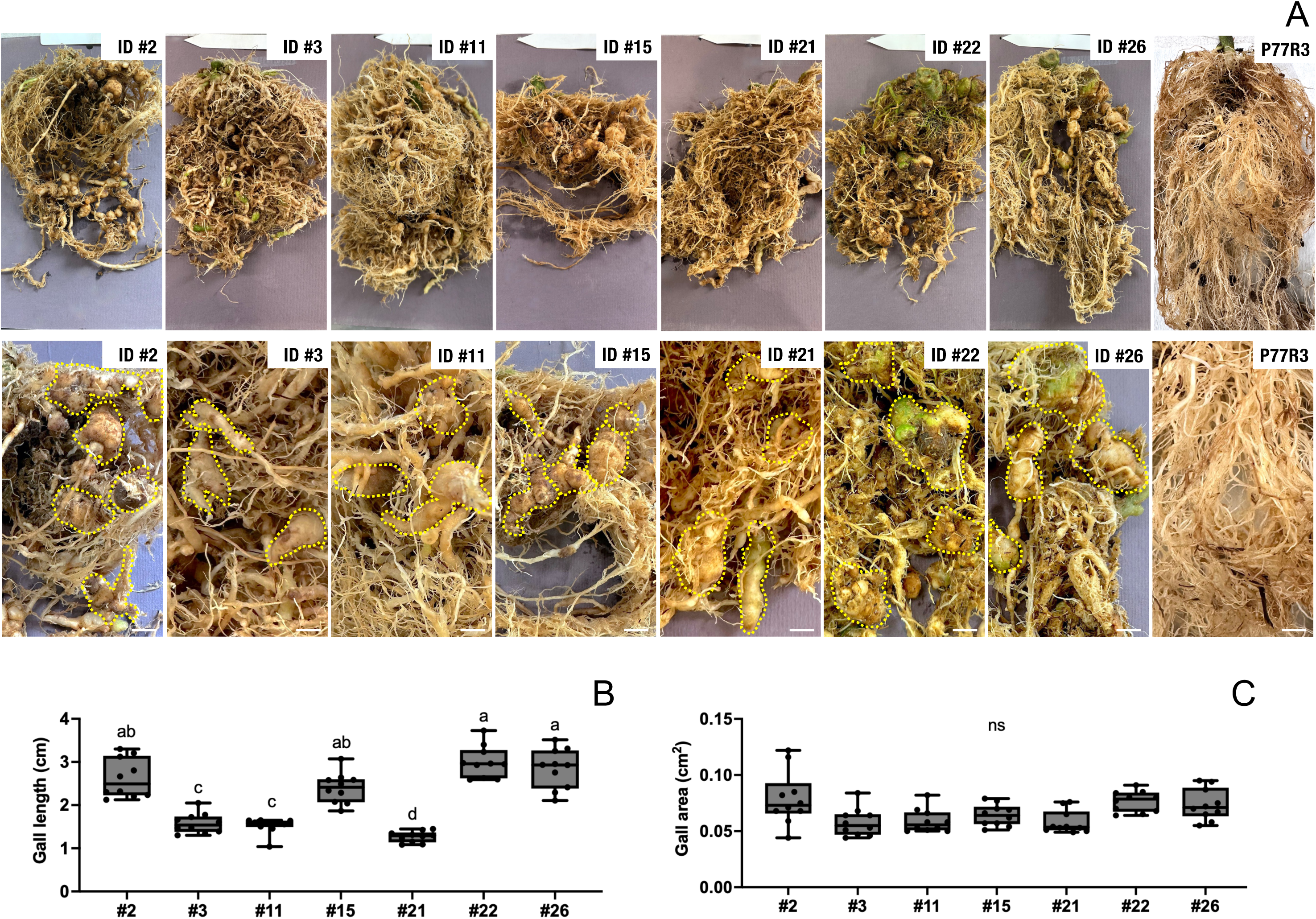
Comparison of galling and root damage measurements among *Meloidogyne* populations. A. Representative images of root systems that had been infected with nematodes from populations #2, 3, 11, 15, 21, 22, and 26 at eight weeks after inoculation. B. *Meloidogyne* populations virulent on *Mi-1* tomato differed significantly in the lengths of the galls they induced on *Mi-1* tomato cv. Celebrity (Kruskal-Wallis test). C. *Meloidogyne* populations virulent on *Mi-1* tomato did not differ significantly in the areas of the galls they induced on *Mi-1* tomato cv. Celebrity (Kruskal-Wallis test). Letters above treatment groups in B and C indicate statistically significant differences, means and standard errors of the mean are shown.

## Discussion

It has been reported that resistance to RKNs in *Mi-1* tomato cultivars is breaking down as different *Meloidogyne* populations evolve mechanisms to evade or suppress (‘break’) resistance mediated by this gene. Resistance-breaking populations have been reported from California and other states within the US as well as from other tomato-growing areas around the globe (Castagnone-Sereno et al., 1993; Hajihassani et al., 2022; Kunwar et al., 2024; Ploeg et al., 2023). In this study, we report a novel resistance-breaking population and identify it as *M. incognita*. Recent studies indicate that *M. incognita* is currently the most common species found in processing tomato agroecosystems across California (Ploeg et al., 2023).

Our study further confirms significant variation in aggressiveness (as indicated by reproductive fitness as a proxy) among *Meloidogyne* populations that are virulent on *Mi-1* tomato. Differences in RF values among such *Meloidogyne* populations have been reported previously (Hajihassani et al., 2022; Ploeg et al., 2023), but our results expand these patterns to a larger group of populations. Given the distinct genetic backgrounds of the populations and the consistent patterns in fitness across generations of these populations, our data suggest that this variation in reproductive ability is at least partially genetic in nature, is heritable, and could thus evolve further. These observations confirm similar findings with resistance-breaking RKN populations on *Mi-1* tomato in other geographical areas (Castagnone-Sereno et al., 1993, 1994). The fact that we observe variation in reproductive ability among *Meloidogyne* populations on a single host genotype—tomato cv. Celebrity with the dominant *Mi-1* gene (Williamson, 1998)— that was infected in a shared, controlled environment, provides further support for the conclusion that variation in this trait is due to genetic variation among the *Meloidogyne* populations rather than genetic or environment-induced differences in host responses. The significant differences in the length of galls induced by the populations provide a third piece of evidence that aggressiveness (as measured by reproductive fitness) has a genetic basis in RKN, since both parasite reproduction and induction of host symptoms are common measures of this trait (Pariaud et al., 2009). Future studies could extend our findings to additional host symptoms related to aggressiveness such as stunting, wilting, and reduced fitness or yield.

Given the evidence that aggressiveness of RKNs has a genetic basis and is heritable, it is possible for variation in this trait to have evolved among *Meloidogyne* populations that occur in different areas of California. It has been proposed previously that populations of pathogens and parasites such as RKNs may evolve higher levels of aggressiveness towards hosts in climates that are better suited for development and reproduction of the pathogen or parasite (Pariaud et al., 2009). In the case of thermophilic species of RKN such as *M. incognita* and *M. javanica*, development and reproduction can be promoted by warmer temperatures throughout the year (Dávila-Negrón and Dickson, 2013). Using latitude as a proxy for geographical variation in year-round temperatures (NOAA NCEI, 2023), we identified a significant correlation between source environments and RF values of the *Meloidogyne* populations when infecting a common host in a common environment. This suggests that populations from lower latitudes may have evolved stronger aggressiveness in the form of higher rates of reproduction following periods of infecting *Mi-1* tomato in environments that are highly conducive to their development and reproduction. It will be interesting to see if the latitudinal gradient in aggressiveness remains apparent when infections take place across a range of temperatures, as temperature changes could potentially influence distinct pathogen or parasite populations differently through genotype-environment interactions (Hargreaves, 2024; Laine, 2008; Pariaud et al., 2009; Price et al., 2004).

We can conclude that *M. incognita* is currently the most common species of RKN virulent on *Mi-1* tomato across Californian processing tomato agroecosystems. We found evidence for the existence of a genetically based latitudinal gradient in aggressiveness (measured here as reproductive fitness) among *Meloidogyne* populations from across California on *Mi-1* tomato, where populations from Southern California tended to be more aggressive than ones from further North. Differences in the length of galls induced by various populations further confirmed a heritable genetic basis for aggressiveness. Our discovery that genetic and phenotypic variation in RKN aggressiveness on *Mi-1* tomato evolved along a latitudinal gradient across California may inform local management practices, including applications of nematicides, across areas that vary in climate. Finally, our findings provide a springboard for further studies on the evolutionary genetics of aggressiveness among *Meloidogyne* populations in different environments. More detailed mechanistic insights into the evolution of this trait among RKN populations could have important implications for ensuring the future productivity of processing tomato cultivation around the globe.

## Acknowledgements

The authors want to thank the National Institute of General Medical Sciences of the National Institutes of Health (grant R35GM151194 to S.C.G.), the National Institute of Food and Agriculture (grant W5186 to S.C.G.), and the University of California Riverside for providing funding for this work.

